# Distinct phosphorylation signals drive acceptor versus self-ubiquitination selection by Parkin

**DOI:** 10.1101/2021.06.01.446590

**Authors:** Karen M. Dunkerley, Anne C. Rintala-Dempsey, Guilia Salzano, Roya Tadayon, Dania Hadi, Kathryn R. Barber, Helen Walden, Gary S. Shaw

## Abstract

The RBR E3 ligase parkin is recruited to the outer mitochondrial membrane (OMM) during oxidative stress where it becomes activated and ubiquitinates numerous proteins. Parkin activation involves binding of a phosphorylated ubiquitin (pUb), followed by phosphorylation of parkin itself, both mediated by the OMM kinase, PINK1. However, targeted mitochondrial proteins have little structural or sequence similarity, with the commonality between substrates being proximity to the OMM. Here, we demonstrate that parkin efficiently ubiquitinates a mitochondrial acceptor pre-ligated to pUb and phosphorylation of parkin triggers autoubiquitination activity. Mitochondrial target proteins, Miro1 or CISD1, tethered to pUb are ubiquitinated by parkin more efficiently than if alone or Ub-tethered and ubiquitin molecules are ligated to acceptor protein lysines and not pUb. Parkin phosphorylation is not required for acceptor-pUb ubiquitination. In fact, only phospho-parkin induced self-ubiquitination and deletion of Ubl or mutation at K211N inhibited self-ubiquitination. We propose divergent parkin mechanisms whereby parkin-mediated ubiquitination of acceptor proteins is driven by binding to pre-existing pUb and subsequent parkin phosphorylation triggers autoubiquitination. This finding is critical for understanding parkin’s role in mitochondrial homeostasis and has implications on targets for therapeutics.

## Introduction

Parkinson’s disease (PD) affects approximately 1% of the world’s population aged 60+ years and is caused by both genetic and environmental factors [1]. Though age of onset and disease severity can vary greatly among patients, there is increasing evidence that implicates mitochondrial dysfunction as a major contributor to neurodegeneration [2]. Genetic forms of PD account for 10% of diagnoses and one such form is autosomal recessive juvenile Parkinson’s disease (ARJP), characterized by onset at 40 years of age or younger. A number of genes have been associated with ARJP but mutations in two genes in particular are found in over 50% of patients: PARK2, which encodes the E3 ubiquitin ligase parkin, and PARK6, which encodes PTEN-induced kinase 1 (PINK1) [3]. Parkin belongs to the RBR E3 ligase family, which contain the characteristic RING1, in-Between-RING, RING2(Rcat) (RBR) domains [4] and utilize a RING/HECT hybrid mechanism of ubiquitin (Ub) transfer [5]. In addition to the RBR domains, parkin has an N-terminal ubiquitin-like (Ubl) domain (30% similarity to ubiquitin) followed by the RING0 domain which is unique to parkin. Under basal conditions, parkin is autoinhibited due to an intramolecular domain associations that inhibit activity [6–8]. Additionally, the catalytic C431 residue in the RING2(Rcat) domain of parkin is partially occluded by an interaction with RING0. These modes of inhibition are relieved by phosphorylation of Ub (pUb) and the Ubl domain in parkin by PINK1 [9–15], and binding of the pUb to parkin [8, 16–18]. Upon oxidative stress, PINK1 is stabilized at the outer mitochondrial membrane (OMM) allowing it to phosphorylate nearby proteins and this triggers localization of parkin to the damaged mitochondrion where it is activated by pUb followed by phosphorylation of the Ubl domain (pParkin) by PINK1 [19–21]. This model indicates that fully activated pParkin ubiquitinates nearby membrane proteins and signals for removal of the damaged mitochondrion [22–24]. Together, parkin and PINK1 work by signaling damaged mitochondrial proteins for degradation and targeting them to the autophagosome. Since substitutions in either of these proteins have detrimental effects on neuronal health, there is a wealth of research that describes how the parkin/PINK1 pathway functions and how defects in the mechanism lead to disease development.

Many groups have worked to identify substrates of parkin including more than 30 potential targets at the mitochondrion alone, especially after oxidative stress. Unbiased techniques like mass spectrometry [22, 25, 26] have identified an extensive network of potential parkin substrates including nuclear, cytoskeletal and mitochondrial proteins following artificial induction of oxidative stress in different cell types. Despite some differences in the cellular treatments used, proteins such as mitofusin 1/2, Miro1, proteins in the TOM complex and CISD1 (MitoNEET) have been identified that are common to the OMM. These OMM proteins have important roles in mitochondrial homeostasis including fusion, transport or axonal movement, so being the early targets of parkin degradation would aid in containing oxidative damage. Close inspection of these substrates does not reveal a common parkin recognition sequence or predict the locations for ubiquitination. So how does parkin select a protein for ubiquitination?

Historically, the activity of parkin has been probed through its autoubiquitination activity where the enzyme was first noted to be inactive in its resting state [6]. Subsequently, increased ubiquitination activity upon recruitment of pUb was noted that is further enhanced by phosphorylation of the parkin Ubl domain [13, 16]. Autoubiquitination of human parkin has been shown to result in multi-ubiquitinated species [27] throughout the protein including K27, K48, K76, K129 and K349 [22, 25, 28]. Further, of the many substrates suggested for parkin, most experiments result in only mono- or di-ubiquitinated products and rarely show the extensive ubiquitination patterns that are observed for autoubiquitination of parkin [8, 16, 29–33]. One of the difficulties encountered is that different methods are frequently used to examine autoubiquitination and ubiquitination of potential substrates that makes comparison and quantifying ubiquitination levels complicated. Nevertheless, in the presence of identified substrates, most experiments show that pParkin is the priority target for ubiquitination and substrates are poorly modified [8, 30, 33, 34]. This observation suggests that substrate ubiquitination may be a result of inefficient association with parkin—a phenomenon that is challenging to explain for a highly regulated enzyme involved in an important part of cellular homeostasis. Immunoprecipitation assays from cell lysates have hinted at an interaction between parkin and some substrates including Miro1 [35, 36] and mitofusin 2 [37] that is enhanced after oxidative stress, however direct interaction experiments or quantification have not been reported. Recently, it has been shown that ubiquitin chains, phosphorylated under oxidative stress conditions, recruit parkin to the OMM [38, 39]. Further, artificial substrates targeted to the OMM and tagged with Ub accelerate parkin-catalyzed ubiquitination upon phosphorylation [39]. These events appear to be dependent on the activity of another accessory E3 ligase, for example MITOL, proposed to seed OMM proteins with ubiquitin that are subsequently phosphorylated to facilitate parkin recruitment [39, 40]. However, *in vivo* it has been difficult to assess the direct ubiquitination of potential OMM substrates based on this model and its relationship to parkin auto-ubiquitination.

Understanding the relationship between parkin activation and substrate targeting is necessary as research proceeds with determining a targetable site on parkin for drug development. Currently, most parkin-based therapeutics are being designed to stimulate parkin ubiquitination by disrupting domain-domain interactions, with regions of interest originating from existing crystal structures [41]. The solved structures acting as models include parkin with all domains present [8, 17], lacking the Ubl domain [7, 42, 43], bound to pUb with and without the Ubl domain [8, 18, 33] and various activated pParkin/pUb forms [44–46]. While these structures have provided details of the specific domain rearrangements upon parkin activation, they only represent snapshots and show no hints of substrate recognition. As previously described, fully activated parkin has a preference for autoubiquitination. Therefore, approaches that rely on targeting an increase in parkin activation may not achieve the necessary therapeutic effect.

In this work we aimed to distinguish the mechanisms used by parkin to select OMM proteins for ubiquitination compared with its self-ubiquitination. We have focused on two OMM proteins, Miro1 and CISD1, known to be ubiquitinated by parkin and created a series of unphosphorylated and phosphorylated substrate-Ub chimeric proteins, or Ub ‘acceptors’, to examine their direct interactions with parkin. Using fluorescent tags that specifically report on the ubiquitination of either an OMM protein or parkin self-ubiquitination we show that pUb alone targets OMM proteins for parkin ubiquitination while phosphorylation of parkin increases its auto-ubiquitination. Collectively, these findings are evidence of divergent mechanisms for parkin ubiquitination activity that show how parkin recognizes OMM proteins needed for mitochondrial clearance under oxidation stress and potentially self-regulates its own degradation.

## Materials & Methods

### Molecular biology of constructs

DNA for the GTPase2 domain of human Miro1 (mG2, residues 402-582) and human CISD1 (residues 32-108) were codon optimized and inserted in a His_6_-expression plasmids (ATUM). The mG2 protein contained 3 point substitutions (V418R, Y470S, L472A) shown to stabilize the monomer form of the protein [34]. For solubilization of mG2, an SMT3 tag was inserted after the His_6_-tag by Restriction-Free (RF-) cloning [47, 48]. Any constructs that required fluorophore labeling, a single Cys residue was inserted by site-directed mutagenesis N-terminal to the protein coding sequence.

The chimeric ‘acceptor-Ub’ species were created by inserting the yeast Ub sequence C-terminal to the substrate, including a Ser-Asn-Ala linker via RF-cloning. mG2_TEV_-Ub was also created by RF-cloning but included the full Tobacco Etch Virus protease (TEV) cleavage site in the inserted DNA sequence. His_6_Ypet_TEV_-Ub was received from Dr. Jiayu Liao (UC Riverside, California). His_6_Ypet_TEV_-Ub was created by inserting the TEVp cleavage site C-terminal to the Ypet fluorophore via a modified RF-cloning protocol.

### Protein Constructs and Purification

All constructs were expressed in *Escherchia coli* BL21(DE3) cells in Luria broth [49], grown at 37 °C until an optical density at 600 nm (OD_600_) of 0.6 was reached. Cells were induced by addition of IPTG were left overnight at 16 °C. All human parkin (full-length, R0RBR, ^77^R0RBR) and Miro1 (mG2, monomeric Miro1 GTPase2 with I459T substitution (mG2^T^) (additional mutation at I459T), mG2-Ub, mG2^T^-Ub, mG2_TEV_-Ub) were encoded as His_6_-Smt3 fusion proteins. For purification, cells were harvested, resuspended in lysis buffer (50 mM Tris, 500 mM NaCl, 0.5 mM triscarboxyethylphosphine (TCEP), 25 mM imidazole, pH 8.0) and lysed using an EmulsiFlex-C5 homogenizer (Avestin). All parkin constructs were purified by Ni^2+^-NTA affinity chromatography by HisTrap FF column on an AKTA FPLC (GE Healthcare) and the affinity tag was cleaved by Ulp1 protease (1:50 protease:protein). After cleavage, the proteins were passed again through the HisTrap FF column and the flow through was collected, concentrated and buffer exchanged by gel filtration column (Superdex75 10/300 GL column, GE Healthcare). Finally, the purified parkin was aliquoted and flash frozen for storage at −80 °C. For all Miro1 constructs, the proteins were purified using Ni^2+^-NTA Sepharose. After batch binding for 1 h at 4 °C, proteins were eluted and rapidly diluted before overnight incubation with Ulp1 protease in 50 mM Tris, 200 mM NaCl, 100 mM Arginine Hydrochloride (ArgHCl), 250 *μ*M TCEP, pH 8.0. The cleaved proteins were then passed through the Ni^2+^-NTA Sepharose and the flowthrough was collected, concentrated and buffer exchanged by gel filtration. The final purified protein was aliquoted and stored at −80 °C.

Yeast Ub and human CISD1 (residues 32-108) were encoded as His_6_-fusion proteins, containing a TEVp site for affinity tag cleavage. His_6_-Ub was loaded onto a Ni^2+^-NTA Sepharose, as above, and eluted. The His_6_ tag was cleaved by adding TEVp (1:50 protease:protein) and the cleaved protein was repassed over the Ni^2+^-NTA Sepharose. The flowthrough was collected, buffer exchange in a gel filtration column, aliquoted and flash frozen at −80 °C. For His_6_-CISD1, a similar purification protocol was used with a few exceptions. The addition of 800 *μ*M FeCl_3_ to the LB media facilitated the expression of the Fe^3+^-bound protein. A buffer containing 20 mM Tris (pH 8.0), 500 mM NaCl, 30 mM imidazole and 250 *μ*M TCEP was used during the cell lysis and Ni^2+^-NTA purification steps. Protein was eluted with 400 mM imidazole and the His_6_-tag was cleaved with TEV protease overnight at 4 °C. The Ypet_TEV_-Ub construct was also purified via Ni^2+^-NTA Sepharose, however the N-terminal His_6_-tag was non cleavable. After the initial affinity column, the Ypet_TEV_-Ub was dialyzed in 50 mM Tris, 200 mM NaCl, 0.25 mM TCEP, pH 7.9 and stored at −80 °C.

Miro1^181-579^ was purified as previously described by Kumar et al. (2015) [8].

P. humanus PINK1 was encoded as a glutathione S-transferase (GST-) fusion and purified as previously described [50]. Briefly, the cells were lysed in PBS buffer, loaded onto a GSTrap FF column (GE Healthcare) and bound protein was eluted with PBS plus 20 mM reduced glutathione (GSH). The purified protein was dialyzed to remove the GSH and finally stored at −80°C for future use.

### Preparation of Phosphorylated Parkin, Ubiquitin and Acceptor-Ub

Purified GST-PINK1 was incubated with full length parkin (1:2), ubiquitin (1:10) or substrate-Ub (1:20) proteins and dialyzed against buffer containing 50 mM Tris, 1.0 mM DTT, 100 mM ArgHCl (mG2-Ub only), 10 mM MgCl2 and 10 mM ATP (pH 7.0) at 25 °C for 16 h. Protein phosphorylation was monitored to completion using Phos-Tag AAL™ (NARD Institute Ltd) SDS–PAGE. Following the reaction, GST-PINK1 was removed using a GSTrap FF column, collecting the flow-through fractions containing the phosphorylated proteins and buffer exchanged by gel filtration. The phospho-proteins were buffer exchanged using a Superdex75 10/300 GL column (GE Healthcare).

### Preparation of mG2^K572^~Ub

Equimolar amounts of mG2 and His_6_-Ub (180 *μ*M) were incubated at 37 °C with E1 (50 nM), UbcH7 (4 *μ*M), pParkin (75 nM) and ATP (10 mM) for up to 2h. To isolate mG2^K572^~Ub, the reaction was purified by Ni^2+^-NTA affinity chromatography and TEVp cleavage, as described above. The final mG2^K572^~Ub was flash frozen and stored at −80 °C.

### Preparation of Fluorescently Labelled Proteins

Ubiquitin, mG2, mG2-Ub, mG2_TEV_-Ub, mG2^K572^~Ub or CISD1-Ub with an N-terminal cysteine (^Cys^protein) (in 25 mM Tris, 100 mM NaCl, 100 mM ArgHCl (mG2 constructs only), 250 μM TCEP, pH 7.4) were expressed and purified as described above. Purified ^Cys^Ub was modified with DyLite 800 maleimide (Thermo Fisher) at a 1:1.5 ratio for 1 hour and quenched with 5 mM DTT. The substrate proteins were modified with a 1:0.8 ratio Alexa Fluor 680 (Thermo Fisher) for 10 min and quenched with 5 mM DTT. Excess DyLight 800 or Alexa Fluor 680 label was removed using a HiLoad Superdex75 gel filtration column. Aliquots of labeled protein were flash frozen and stored at −80 °C until needed.

### Isothermal Titration Calorimetry

All calorimetry experiments were performed using a NanoITC (TA Instruments) at 18 °C (mG2^T^-Ub) or 25 °C (CISD1-Ub). All experiments were completed 2 times using freshly prepared proteins in 25 mM HEPES, 100 mM NaCl and 250 *μ*M TCEP at pH 7.5. The concentrations of each protein in the experiments were as follows: 20 *μ*M parkin titrated with 100 *μ*M mG2^T^-Ub, 84 *μ*M mG2^T^-pUb, 200 *μ*M CISD1, 212 *μ*M CISD1-Ub and 106 *μ*M CISD1-pUb. Data were analysed using single-site binding models using NanoAnalyze software (TA Instruments).

### Microscale Thermophoresis

His_6_-Smt3-Parkin constructs were labelled with Monolith His-Tag Labeling Kit RED-tris-NTA 2nd Generation (Cat#MO-L018) according to the supplied labelling protocol. The unlabelled Miro1181-579 was used as the ligand. A titration series of up to 16 dilutions was generated as a 1:1 dilution of the previous sample, starting from a ligand concentration of 50 *μ*M. All the experiments were carried out in PBS-T buffer (PBS 1x, 0.05% Tween 20). The MST experiments were performed on a NanoTemper Monolith NT.115. Measurements were performed at 25 °C using 60% LED and 40% MST power, with laser off/on times of 5 and 30 sec, respectively. Binding affinities were determined by using a 1:1 binding interaction, and data were fitted with a Hill equation 1 (Hill coefficient = 1):

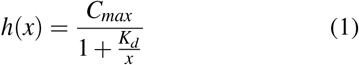

where *C_max_* is the fluorescence difference the bound and unbound state, and *K_d_* is the dissociation constant. The fit was performed using least-square minimization method in GNUPLOT software.

### Analytical Ultracentrifugation

Sedimentation velocity experiments were conducted using a Beckman XL-A analytical ultracentrifuge equipped with an An60Ti rotor. Double sector cells (1.2 cm) with quartz windows were filled with 380 *μ*L sample and 400 *μ*L reference buffer. All data were collected in 50 mM Tris, 200 mM NaCl, 0.5 mM TCEP pH 8.0 at 18°C (mG2-Ub) or 20 mM Tris, 100 mM NaCl, 1 mM DTT pH 8.0 at 20°C (CISD1-Ub). All experiments were performed at 45,000 rpm. Cells were scanned at equal intervals, 45 scans total, and sedimentation was monitored by absorbance at 250 nm (mG2-Ub) or 280 nm (CISD1-Ub). Data were processed by c(s) distribution analysis in SEDFIT v15.01, accounting for equilibration and rotor acceleration times. For parkin, mG2-Ub and CISD1-Ub, partial specific volume 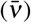 was 0.719 mL g^−1^, 0.739 mL·g^−1^, 0.7377 mL·g^−1^, respectively, viscosity (*h*) was 0.01033 Poise, and density (*r*) was 1.0079 g/mL. Sedimentation coefficients observed in buffer were corrected for 20 °C in water (S_20,w_), and frictional coefficients (f/f_0_) were calculated from S_20,w_ and known molecular weights of all species. All data were fit to an RMSD <0.005 in SEDFIT.

### Parkin Ubiquitination Assays

Most reactions were completed with 4 *μ*M Ub, 1 *μ*M ^800^Ub, 0-2 *μ*M pUb, 1 *μ*M (p)Parkin, 2 μM substrate (mG2, mG2-Ub, mG2_TEV_-Ub), 0.5 *μ*M UbcH7, 5 mM MgATP, 50 mM HEPES, pH 7.5. For CISD1-(p)Ub reactions, the reactions contained 1 *μ*M UbcH7, 1 *μ*M parkin, 1 *μ*M CISD1-Ub, 1 *μ*M pUb and 20 *μ*M Ub. All assays were initiated by adding 0.1 *μ*M Uba1, reacted at 37 °C and quenched at desired time points using 3x SDS sample buffer and 1 *μ*L of 1M DTT. Gradient gels (4-12% Bis-Tris Plus, Thermo Fisher Scientific) were used with MES running buffer (250 mM MES, 250 mM Tris, 0.5% SDS, 5 mM EDTA, pH 7.3). Gels were scanned by Odyssey Imaging system (LiCor) and fluorescence intensity was measured at 700 nm and 800 nm.

### TEV-cleavage Ubiquitination Assays

The base ubiquitination assays in these reactions were conducted as above. FormG2_TEV_-pUb, the reaction contained 0.5 *μ*M UbcH7, 0.5 *μ*M (p)Parkin, 4.25 *μ*M Ub, 0.75 *μ*M ^800^Ub, 1.7 *μ*M mG2_TEV_-pUb, 0.3 *μ*M ^680^mG2_TEV_-pUb and were initiated by 0.1 *μ*M Uba1. After 15 minutes, ubiquitination activity was halted with 2 *μ*L of 1 mg/mL apyrase and 1.25 *μ*M EDTA. Finally, 2 *μ*L of 2.5 mg/mL TEVp was added and the cleavage progressed for 60 min. All samples were quenched with 3xSDS sample buffer and run on 10% Bis-Tris gels in MES running buffer.

For Ypet_TEV_-pUb, the reactions contained 1 *μ*M UbcH7, 2 *μ*M (p)Parkin, 40 *μ*M Ub, 10 *μ*M Ypet_TEV_-pUb and were initiated by 0.05 *μ*M Uba1. After 60 min, the reactions were incubated with 100 *μ*L of Ni^2+^-NTA Sepharose (Qiagen) in a spin column to trap the Ypet_TEV_-Ub Ubn and remove other reaction components. The column was washed twice with 50 mM HEPES, 50 mM NaCl, pH 7.0. Cleavage was induced by adding 5 *μ*L of 2.5 mg/mL TEVp directly to the immobilized Ypet_TEV_-pUb in 165 *μ*L 50 mM HEPES, 50 mM NaCl, pH 7.0. The cleavage product was collected and any remaining YPet~Ub_n_ was eluted with 150 *μ*L elution buffer. All samples were quenched with 3xSDS sample buffer and run on 10% Bis-Tris gels in MES running buffer. YPet fluorescence was visualized on a BluPAD LED Transilluminator (FroggaBio) and all proteins by Coomassie staining.

## Results

### Parkin auto-ubiquitination exceeds target protein ubiquitination

Parkin autoubiquitination is most commonly used as a determinant of activity and typically shows that parkin is activated by binding of a pUb molecule with enhanced ubiquitination observed following PINK1 phosphorylation of the parkin Ubl domain. We sought to identify a method where we could monitor autoubiquitination and target protein ubiquitination simultaneously. This would allow us to distinguish the mechanisms used by parkin for these processes and gauge the relative efficiencies of both. We used two known parkin target proteins: Miro1 and CISD1 (Fig. 1A) that are both located on the OMM and have been shown in multiple studies to be ubiquitinated by parkin under oxidative stress conditions [24, 30, 31, 33–36, 51]. Miro1 is a multi-domain GTPase anchored to the OMM at its C-terminus. Previous studies have indicated K572 in the GTPase2 domain of Miro1 as the primary site of parkin ubiquitination [22, 30, 34, 52] so we created a minimal protein construct, mG2, that contained only the C-terminal GTPase2 domain of human Miro1 (residues 411-580) (Fig. 1B). Likewise, the redox protein CISD1 (MitoNEET) lacking its transmembrane domain (Fig. 1B) (residues 32-108), was created that contained all identified parkin-mediated ubiquitination sites. These proteins were fluorescently tagged with Alexa Fluor 680 at their N-termini while Ub was modified with a DyLight 800 fluorophore (Fig. 1B). This allowed ubiquitination of Miro1 or CISD1 (red fluorescence) to be distinguished from parkin auto-ubiquitination (green fluorescence) in a single experiment using simultaneous fluorescence detection. Under typical *in vitro* ubiquitination assay conditions parkin autoubiquitination, indicated by green fluorescence, is robust in the presence of pUb and PINK1-phosphorylated parkin (pParkin) (Fig. 1C&D, lanes 1-5), consistent with many other studies [8, 16, 17]. Upon addition of a target protein, either mG2 or CISD1, parkin autoubiquitination remains efficient. However, little change in the red fluorescent band occurs indicating that ubiquitination of mG2 or CISD1 is minimal under identical conditions (Fig. 1C&D, lanes 6-10). These data show that the efficiency of parkin autoubiquitination far exceeds that for either Miro1 or CISD1.

**Figure 1:**
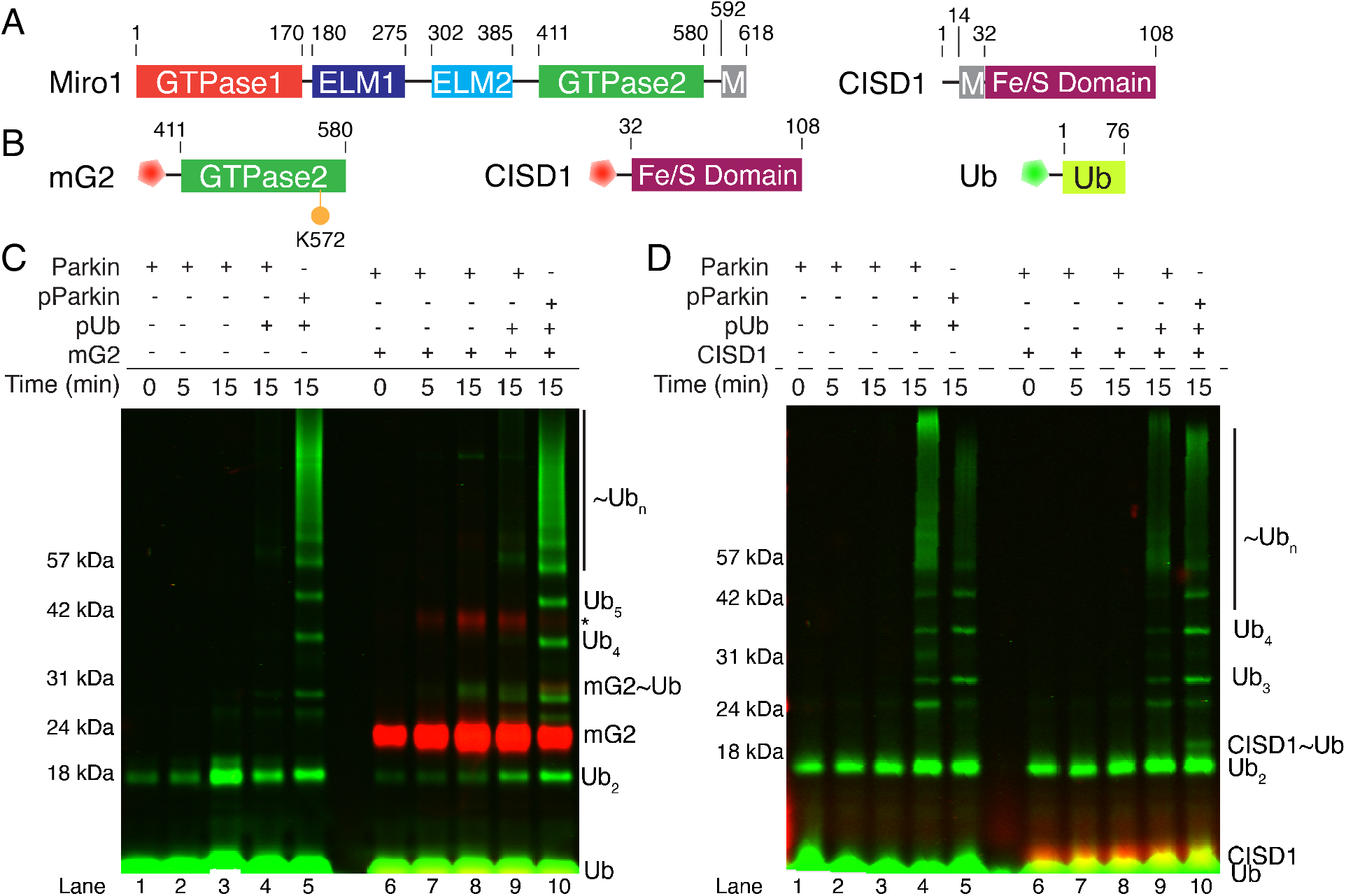
Parkin does not ubiquitinate an isolated substrate, mG2 or CISD1. (*A*) Schematic diagram showing the domains of human Miro1 and human CISD1. (*B*) Schematic diagram showing the minimal target constructs mG2 and CISD1 used in this study. Also indicated are the N-terminal fluorescent labels for mG2, CISD1 (Alexa Fluor 680, red pentagon) and Ub (DyLight 800, green pentagon). (*C*)&(*D*) Parkin autoubiquitination and minimal substrate ubiquitination were monitored by fluorescently labeled ^800^Ub (green) and ^680^mG2 or ^680^CISD1 (red). Reactions were initiated by addition of Uba1 (E1) and quenched at the required timepoints with 3x sample buffer and DTT.

### Phospho-ubiquitin is a parkin substrate that anchors acceptor/target proteins

Parkin is a hybrid RING-HECT E3 ligase which acts as a scaffold for the E2~Ub conjugate [44] and a protein targeted for ubiquitination [44]. Biologically, this mechanism requires recruitment of parkin to the OMM and physical interaction between parkin and proteins such as Miro1 and CISD1 to facilitate ubiquitin transfer. However, limited data is available that shows direct interaction between the proteins. Recruitment of parkin to the OMM has been well-established [19–21]. To test this *in vitro*, we measured the direct interactions between purified parkin and either a soluble construct of Miro1 containing the EF1-EF2-G2 domains (residues 181-579, Miro1^181-579^) or CISD1 using either microscale thermophoresis or isothermal titration calorimetry (Fig. S1). Using protein concentrations much higher than anticipated in cells we were unable to detect any direct interactions between parkin and either of these proteins. Based on the concentrations used in these experiments these data indicate that the interactions of CISD1 or Miro1^181-579^ with parkin are extremely poor with estimated dissociation constants (dissociation constant (K_d_) greater than 100 *μ*M. This indicates that OMM proteins such as Miro1 and CISD1 are not genuine substrates that interact directly with parkin.

It has been proposed that pre-ubiquitination of an OMM protein by an endogenous mitochondrial E3 ligase [39] and subsequent phosphorylation of Ub by PINK1 accelerates parkin recruitment to damaged mitochondria [38] and enhances ubiquitination of artificial substrates such as maltose-binding protein. Such a mechanism would also account for immunoprecipitation studies which show that parkin can be pulled down from a cell lysate by Miro1 [35, 36]. To identify how “pre-ubiquitination” controls these events with OMM proteins, we created two chimeric proteins where the N-terminus of Ub was fused via a Ser-Asn-Ala linker to the C-termini of mG2 and CISD1, creating two pseudo-ubiquitinated target proteins mG2-Ub and CISD1-Ub (Fig. 2A). We then used ITC to measure the strengths of interaction with each of these chimeric proteins with parkin. In both cases, a measurable interaction could not be observed (Fig. 2B) indicating that Ub attached to either mG2 or CISD1 is incapable of promoting the association of the free protein with parkin. We verified these observations using sedimentation velocity analytical ultracentrifugation which shows complexes were not formed between parkin and either mG2-Ub or CISD1-Ub (Fig. S2A). Each of these chimeras could be efficiently phosphorylated at S65 of Ub by PINK1 (mG2-pUb or CISD1-pUb) to test whether the phosphorylated chimera might have an altered interaction with parkin. When titrated into parkin, tight binding is observed with measured dissociation constants (K_d_) of 894 nM and 428 nM, respectively (Fig. 2C) for mG2-pUb or CISD1-pUb. In both cases the endothermic heat change and dissociation constants are similar to ITC measurements for the isolated pUb interaction with parkin [8, 17, 51] suggesting the pUb interaction for the chimeric proteins is as for pUb alone. Further, since the dissociation constants for mG2-pUb or CISD1-pUb were not enhanced compared to pUb alone this suggests that additional interactions do not exist between the target mG2 or CISD1 proteins and parkin that would be expected to increase the observed affinities. These interactions were validated using sedimentation velocity experiments that show mixtures of parkin with either mG2-pUb or CISD1-pUb yielded new signals with higher sedimentation coefficients (Fig. S2) than parkin alone, indicative of tight complex formation for both phosphorylated chimeras with parkin. Our results indicate that phospho-ubiquitin is the primary determinant that targets the interaction of an OMM protein with parkin.

**Figure 2:**
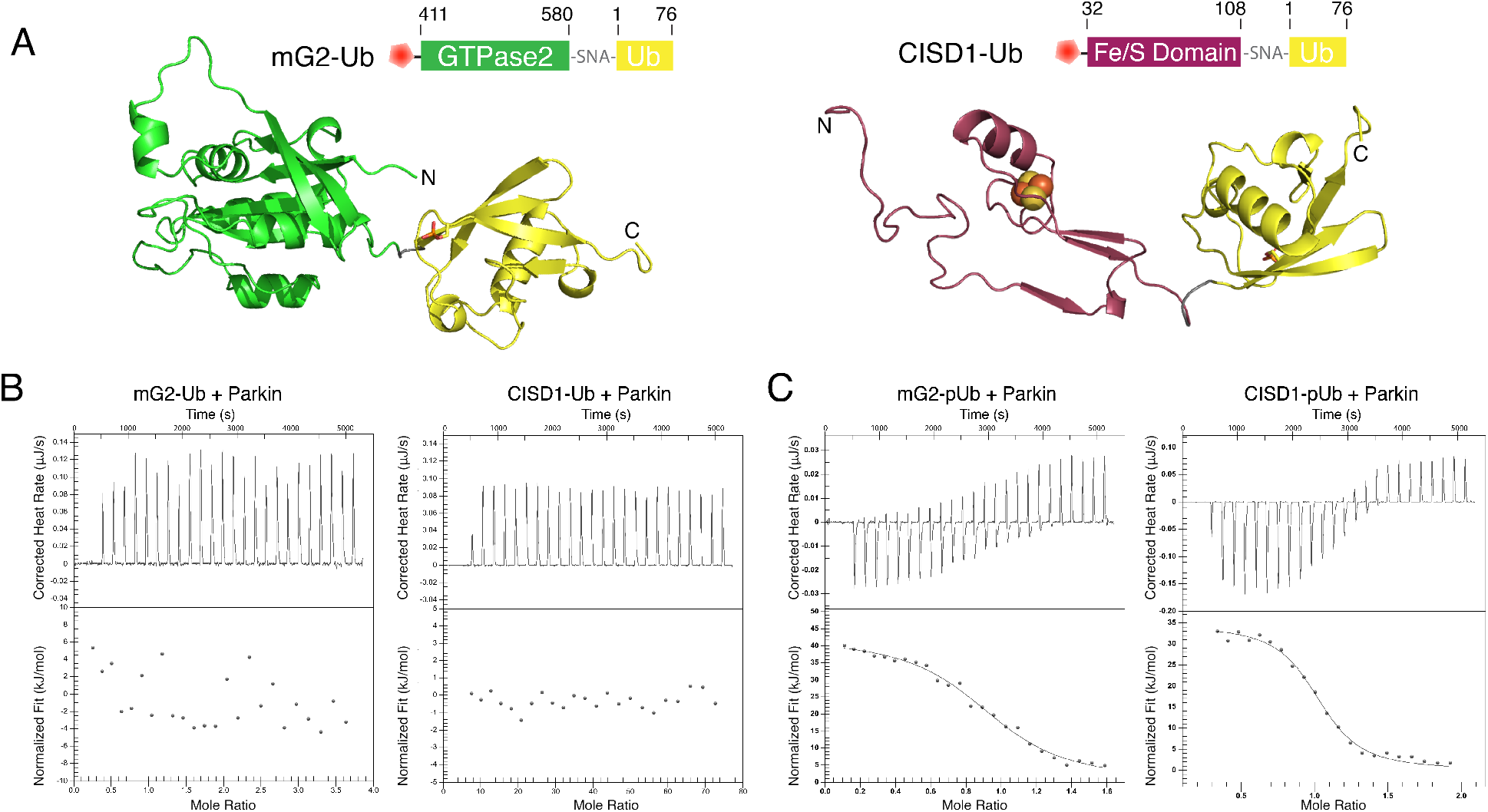
Parkin interacts with a substrate-pUb. (*A*) Schematics and models of chimeric constructs mG2-Ub (modified PDB: 5KSO and 1UBQ) and CISD1-Ub (modified PDB: 3REE and 1UBQ). Structures were modified using Chimera and Modeller. (*B*)&(*C*) Isothermal titration calorimetry of parkin (20 *μ*M) titrated with mG2-Ub (100 *μ*M), CISD1-Ub (212 *μ*M), mG2-pUb (84 *μ*M) or CISD1-pUb (106 *μ*M).

We hypothesized that the pUb substrate of the mG2-pUb or CISD1-pUb species plays a critical role in anchoring and positioning the attached protein (an “acceptor”) for efficient ubiquitination by parkin. To test this idea, we conducted parkin-mediated ubiquitination assays using the mG2-Ub and CISD1-Ub chimeras. In the absence of phosphorylation, ubiquitination of mG2-Ub (Fig. 3A, lanes 1-5) or CISD1-Ub (Fig 3B, lanes 1-5) is very inefficient as shown by the lack of red-fluorescent higher molecular weight bands in the gel. On the other hand, a high degree of autoubiquitination is noted for fully-activated pParkin in complex with free pUb shown by the green fluorescent bands. The lack of acceptor ubiquitination is consistent with the poor complex formation between parkin and either mG2 or CISD1 shown in Fig. 2B (also Fig. S1, S2A). In contrast, phosphorylation of mG2-Ub leads to robust ubiquitination of the chimeric protein by parkin, driven by the pUb substrate, as denoted by the red fluorescence spaced at regular molecular weight intervals (Fig 3, lanes 6-9). Though not as striking, more efficient ubiquitination of CISD1-pUb is evident in the intensity decrease of the unmodified CISD1-pUb band (Fig. 3B, lane 8). These observation are more pronounced when the red and green fluorescence are split (Fig. 3, centre and right, respectively). These more clearly show the extent of acceptor-pUb~Ub_n_ by parkin (Fig. 3, red lane 9) with virtually no self-ubiquitination (Fig. 3, green lane 9). Phosphorylation of parkin shifts the distribution of acceptor-pUb~Ub_n_ to higher MW species and increases parkin self-ubiquitination and free Ub-chain formation (Fig. 3A, red and green lanes 10-11).

**Figure 3:**
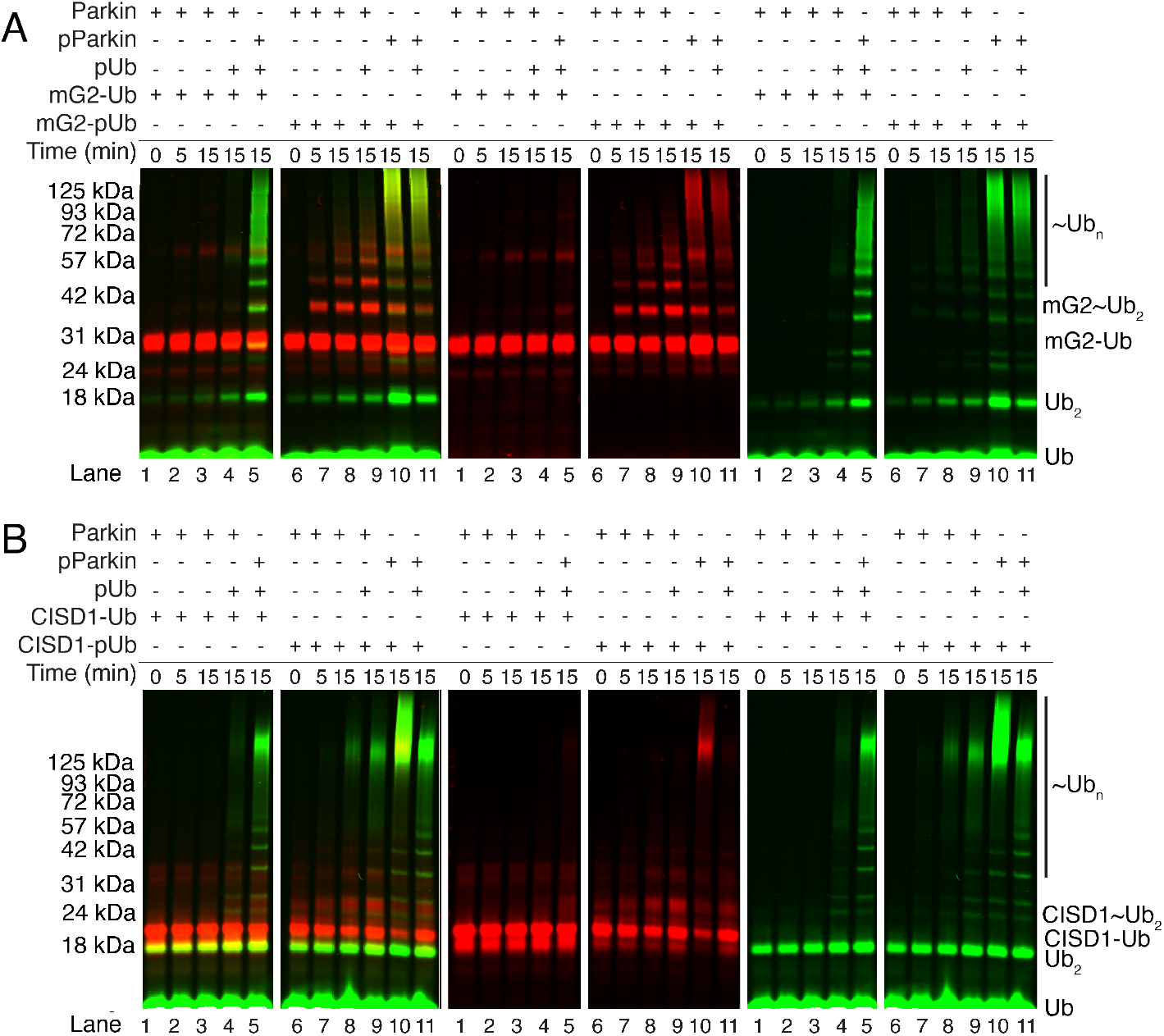
Parkin poly-ubiquitinates the acceptor-pUb. Parkin autoubiquitination of (*A*) mG2-(p)Ub or (*B*) CISD1-(p)Ub ubiquitination were monitored by fluorescently labeled ^800^Ub (green) and ^680^mG2-Ub or ^680^mG2-pUb (red). Reactions were initiated by addition of Uba1 (E1) and quenched at the required timepoints with 3x sample buffer and DTT.

### Parkin uses divergent mechanisms for substrate ubiquitination and autoubiquitination

Our acceptor-pUb chimeras show that anchoring an OMM protein to parkin via a linked -pUb facilitates ubiquitination. This occurs in the absence of PINK1 phosphorylation of parkin, suggesting this step is not essential for acceptor ubiquitination by parkin. Upon phosphorylation of the Ubl domain in parkin, and in the presence of a pUb chimera we observed enhanced green fluorescence from a mixture of ubiquitinated chimera-pUb and pParkin~Ub_n_ species (Fig. 3, lane 10). These observations may indicate that phosphorylation of the Ubl domain acts as a switch to direct substrate ubiquitination or parkin autoubiquitination.

In the presence of pUb binding to parkin, it has been shown that phosphorylation of the Ubl domain weakens its interaction with its RING1 binding site [8, 50] while crystal structures lacking the RING2(Rcat) domain show the phosphorylated Ubl domain has the potential to re-bind to a basic region of the RING0 domain (K161, K163, K211) [45, 46]. Since our data have demonstrated that phosphorylation of parkin is not required for ubiquitination of either mG2-pUb or CISD1-pUb, we created two truncated parkin proteins, lacking either the Ubl-linker domains (R0RBR, residues 141-465) or only the Ubl domain (^77^R0RBR, residues 77-465) to test Ubl-independent acceptor ubiquitination. In the absence of an acceptor both R0RBR and ^77^R0RBR proteins had weaker autoubiquitination activity compared to full-length parkin, even in the presence of pUb (Fig. 4A), in agreement with previous data that shows the phosphorylated Ubl domain is needed for full parkin autoubiquitination activity [7, 8].

**Figure 4:**
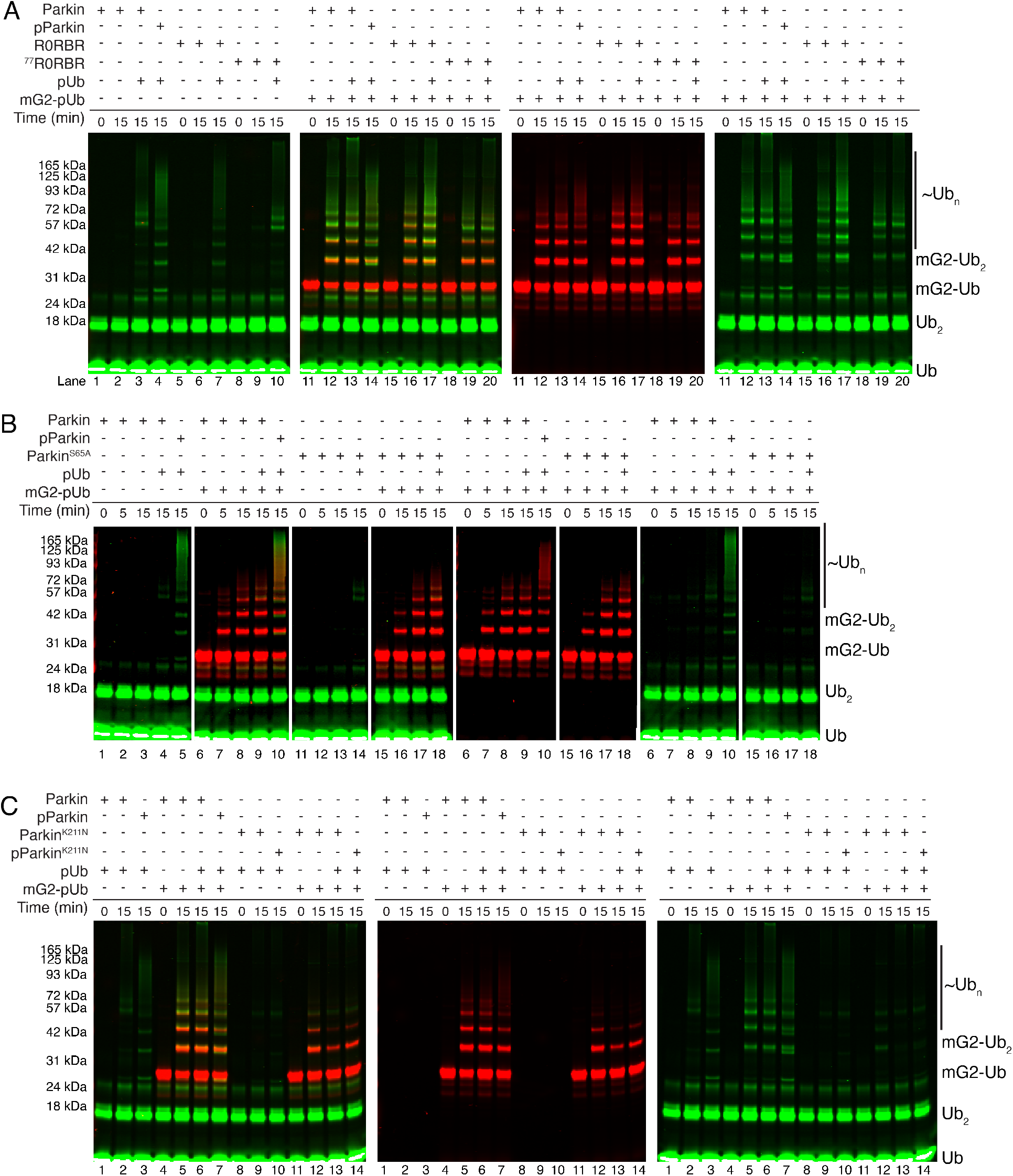
The Ubl domain is not required for mG2-pUb poly-ubiquitination. Ubiquitination assay using fluorescently labeled ^800^Ub (green) and ^680^mG2-pUb (red) to test the activity of (*A*) two truncated parkin’s, R0RBR and ^77^R0RBR, substituted parkin^S65A^ and (*C*) parkin^K211N^. Reactions were initiated by addition of Uba1 (E1) and quenched at the required timepoints with 3x sample buffer and DTT. Images are shown of combined and split red and green fluorescence.

Remarkably, introducing mG2-pUb leads to significant ubiquitination increases by both R0RBR and ^77^R0RBR constructs. As noted from the red fluorescence, much of the ubiquitination occurs on the mG2-pUb protein. Additionally, a non-phosphorylatable parkin, parkin^S65A^, shows ubiquitination of the acceptor-pUb nearly identical to that observed for wild-type parkin. Concurrently, autoubiquitination by parkin^S65A^ is completely eliminated (Fig. 4B). The absence of the pUbl domain in these experiments shows that unlike the mechanism suggested by recent crystal structures, the phosphorylated Ubl domain may not necessary to facilitate acceptor ubiquitination. To test this idea, an early-onset PD substitution in the RING0 domain, K211N, shown to decrease autoubiquitination [20, 53] presumably from a disruption of the more recently observed pUbl-binding site [45, 46]. As expected, parkin^K211N^ has little observable autoubiquitination activity upon phosphorylation of the Ubl domain or in the presence of free pUb (Fig. 4C, left). In contrast, upon addition of the mG2-pUb acceptor, the results were markedly different. For both parkin^K211N^ and pParkin^K211N^, we observed significant ubiquitination of the mG2-pUb species even though the pUbl binding site has been eliminated (Fig. 4C, right). We also observed the near absence of parkin^K211N^ autoubiquitination.

The ubiquitination of the chimeric acceptors was specific for the presence of the pUb anchor since chimeras where parkin’s N-terminal Ubl domain was tethered to mG2 in place of Ub could not drive ubiquitination onto the acceptor, even upon phosphorylation (Fig. 5). The mG2-pUbl chimera failed to activate both acceptor- and autoubiquitination with all parkin variants, including those lacking the Ubl domain (Fig. 5B). These results indicate that OMM target protein ubiquitination is dependent on its association with parkin via an attached pUb substrate while phosphorylation of the Ubl domain further activates parkin and favors autoubiquitination.

**Figure 5:**
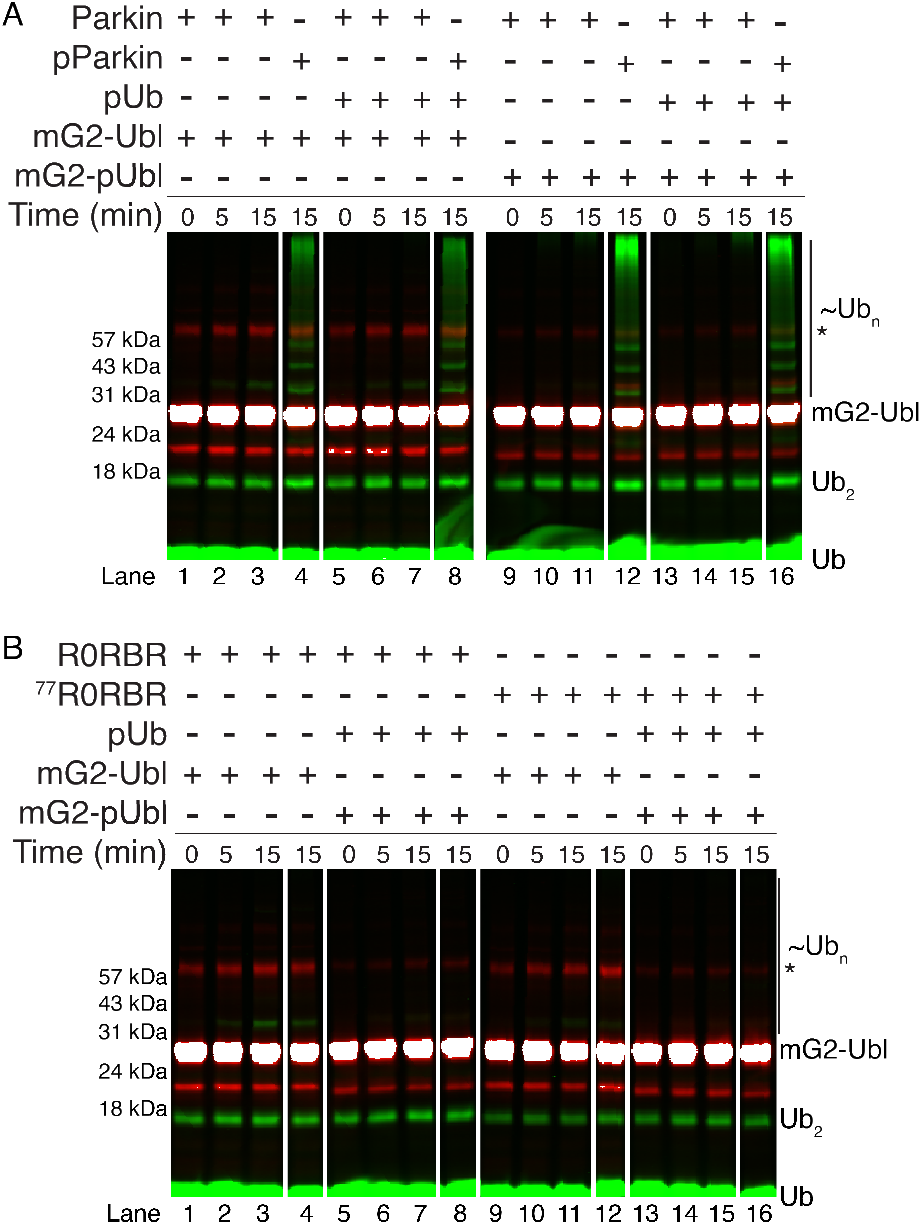
A linked acceptor-pUbl cannot induce ubiquitination by parkin or DUbl parkin. Ubiquitination assays using mG2-Ubl and parkin, pParkin amd two DUbl parkin constructs, R0RBR and ^77^R0RBR. Reactions were initiated by addition of Uba1 (E1) and quenched at the required timepoints with 3x sample buffer and DTT.

We considered the possibility that the C-terminal -GG-sequence in the Ub moieties of each of the chimeric proteins could be used to promote ubiquitin chain formation, rather than free Ub used in the reactions. To test this, we synthesized fluorescently-labeled mG2 that was mono-ubiquitinated at K572 (mG2^K572^~Ub), phosphorylated it using PINK1 and conducted similar ubiquitination assays (Fig. 6, S3). Under these conditions where the conjugated C-terminus of Ub is blocked, we observed efficient poly-ubiquitination of mG2^K572^~pUb. In contrast, the unphosphorylated conjugate mG2^K572^~Ub was poorly ubiquitinated (Fig. 6). In addition, there is minimal improvement in the ubiquitination of mG2^K572^~pUb upon phosphorylation of parkin. Most striking is the robust ubiquitination of the acceptor-pUb by unphosphorylated parkin, with a notable lack of autoubiquitination or high molecular weight bands. Together, these results indicate that the pUb substrate linked to an acceptor is a requirement for acceptor ubiquitination by parkin and that phosphorylation of parkin is not a pre-requisite for this step.

**Figure 6:**
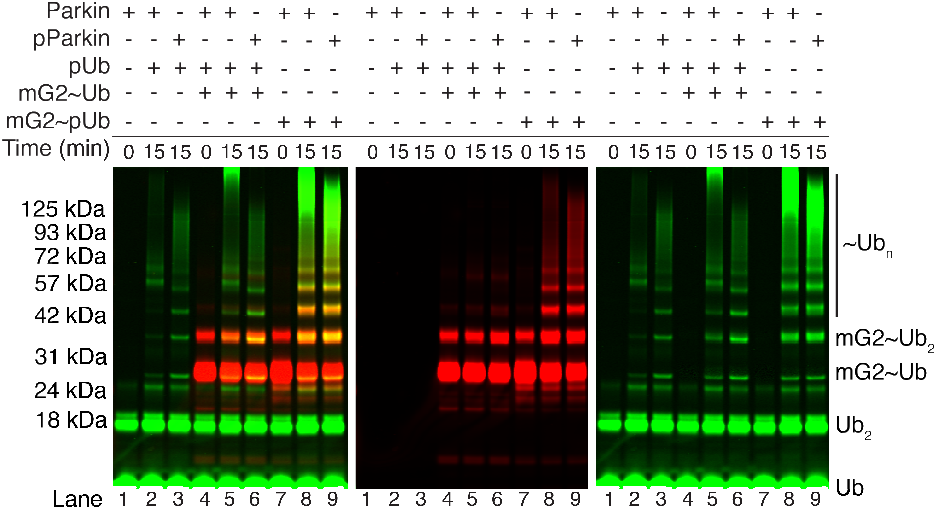
Parkin efficiently multi-ubiquitinates a truly mono-ubiquitinated substrate. Parkin autoubiquitination and mG2^K572^~Ub ubiquitination were monitored by fluorescently labeled ^800^Ub (green) and ^680^mG2^K572^~Ub or ^680^mG2^K572^~pUb (red). Reactions were initiated by addition of Uba1 (E1) and quenched at the required timepoints with 3x sample buffer and DTT.

### Parkin’s phospho-ubiquitin binding site is used by an acceptor-pUb

We used ubiquitination assays to confirm that mG2-pUb binds to parkin in a similar manner as observed for pUb alone. In the absence of chimeric protein, higher pUb concentrations increase parkin autoubiquitination (Fig. 7, lanes 2-5). While pParkin autoubiquitination required the presence of pUb, an observable decrease in activity occurs above stoichiometric amounts of pUb (Fig 7, lanes 6-9). In the presence of mG2^K572^~pUb, both parkin and pParkin show more robust polyubiquitination compared to pUb alone. In both cases, increasing additions of pUb leads to an observable decrease in ubiquitination activity (Fig 7, lanes 11-18). This result suggests that pUb and mG2-pUb compete for the same binding site between the RING0 and RING1 domains where a basic patch is formed by K151, H302 and R305. Substitution at one of these residues, parkin^H302A^, eliminates the ubiquitination of mG2-pUb and significantly reduced pParkin^H302A^ autoubiquitination (Fig 8). Together these observations support our findings that creation of a pUb substrate on an OMM acceptor protein is needed for the acceptor to associate with parkin at the RING0/RING1 cleft and enable ubiquitination of the acceptor-pUb target complex.

**Figure 7:**
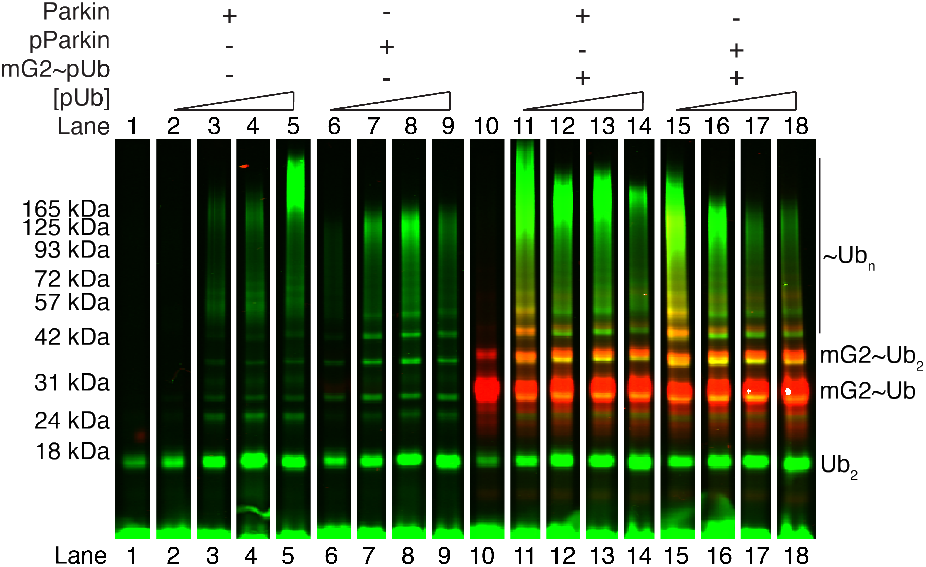
Free pUb and mG2^K572^~pUb compete for the same site on parkin. Ubiquitination assays done in the presence of increasing concentrations of free pUb. Lanes 1 and 10 are time = 0 min. All other lanes are after 15 min of reaction. Ubiquitination was monitored by fluorescently labeled ^800^Ub (green) and ^680^mG2^K572^~pUb (red). Reactions were initiated by addition of Uba1 (E1) and quenched at the required timepoints with 3x sample buffer and DTT.

**Figure 8:**
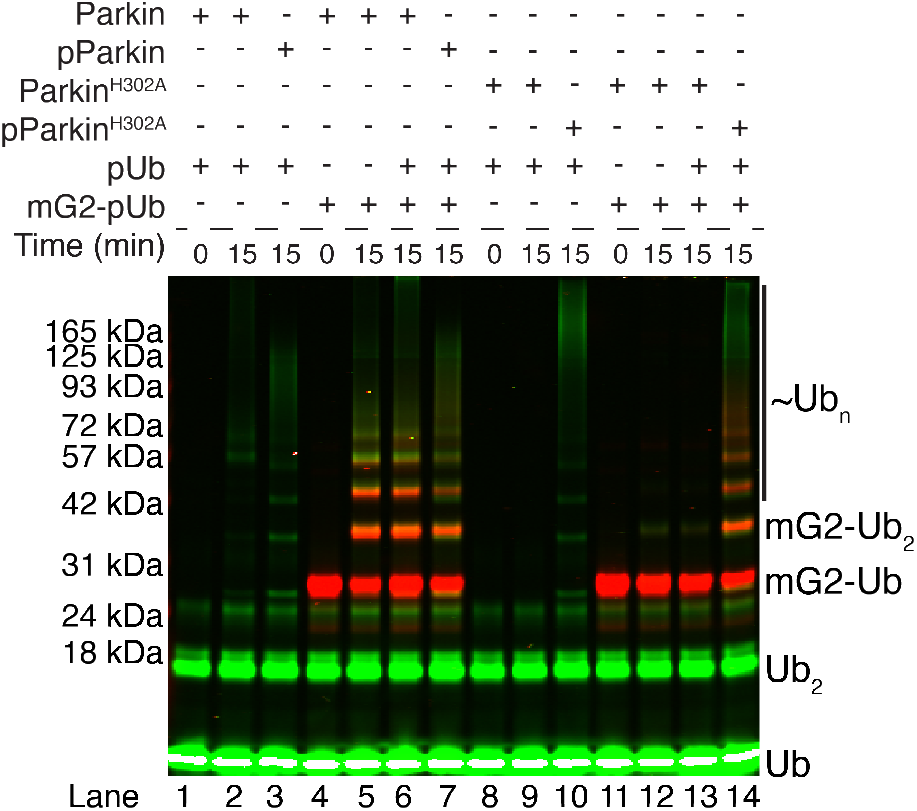
Free pUb and mG2^K572^~pUb bind the same site on parkin. Ubiquitination assays using substituted parkin^H302A^. Activity was monitored by fluorescently labeled ^800^Ub (green) and ^680^mG2-pUb (red). Reactions were initiated by addition of Uba1 (E1) and quenched at the required timepoints with 3x sample buffer and DTT.

### Acceptors are ubiquitinated, not phospho-ubiquitin

In our system we have shown that phosphorylated Ub linked to the OMM proteins Miro1 and CISD1 stimulates parkin ubiquitination of the chimeric proteins. It is possible that either the mG2/CISD1 or pUb components in the chimeras could be ubiquitinated in our assays, leading to potentially different biological outcomes. For example, ubiquitination of the OMM proteins would provide a signal for mitophagy, while ubiquitination of the pUb protein may be an indirect result of the chimeric protein and have little relationship to mitochondrial removal. To identify whether the OMM component of an acceptor-pUb is ubiquitinated, we first inserted a TEV protease cleavage site (VENLYFQ^SN) between the fluorescently-labeled mG2 and pUb moieties that would allow proteolytic cleavage of the chimeric protein, mG2_TEV_-Ub (Fig. 9A).

**Figure 9:**
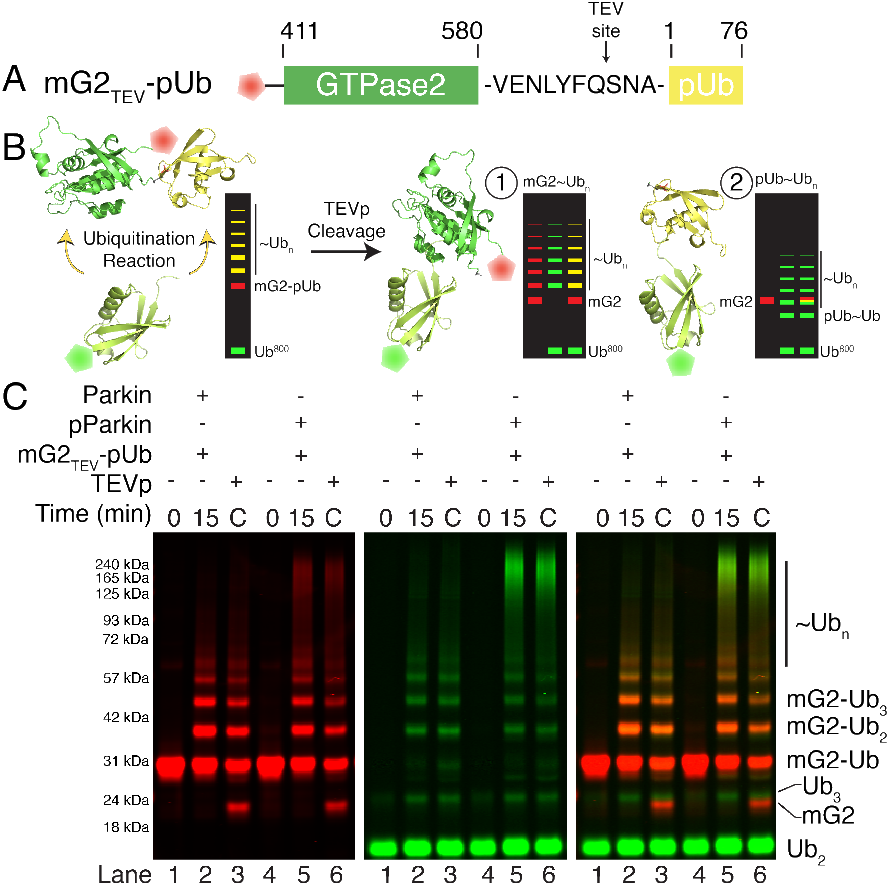
Parkin ubiquitinates the substrate and not the pUb. (*A*) Schematic diagram of the TEV cleavable mG2_TEV_-Ub chimera. (*B*) Model structures (modified PDB: 5KSO and 1UBQ) and results demonstrating the possible results after ubiquitination and cleavage of mG2_TEV_-pUb. C Ubiquitination assay using fluorescently labeled ^800^Ub (green) and ^680^mG2_TEV_--pUb (red). The reactions were left for 15 min before adding apyrase (0.05 mg/mL) and EDTA (1.25 *μ*M) to stop parkin activity and TEV to cleave the ubiquitinated substrate.

Following ubiquitination and subsequent TEV cleavage, two outcomes are possible with respect to the red fluorescent mG2 protein (Fig. 9B). In the first scenario, ubiquitination on the mG2 would yield a red-fluorescent multi-ubiquitin ladder that has been shifted on a gel by the mass of the single phospho-ubiquitin tag that has been cleaved. Alternatively, if ubiquitination occurs on the pUb moiety, a collapse of the red fluorescent ladder to a single mG2 band and shifting of the green ubiquitin ladder to a lower position on the gel by the mG2 mass should be observed. Ubiquitination reactions showed robust modification as denoted by the red fluorescence on themG2_TEV_-pUb protein (Fig. 9C). Upon TEV cleavage, it was noted that a new red fluorescent band corresponding to non-ubiquitinated mG2 appears and the multi-ubiquitinated red ladder from mG2 remains. Further, there is little change in the positions or intensities of the bands for Ub_2_ or Ub_3_ (green fluorescence). These observations indicate that parkin-mediated ubiquitination ofmG2_TEV_-pUb is directed to the acceptor protein and that the pUb substrate acts as an anchoring module that positions the OMM protein for ubiquitination but is not ubiquitinated itself.

To further confirm this, we tested parkin-mediated ubiquitination of a non-biological acceptor *in vitro*. We created a Ypet_TEV_-Ub chimera that has a TEV site for proteolytic cleavage and N-terminal His6-tag for purification following ubiquitination. A similar approach has been used successfully in vivo using an in-frame GFP-Ub fusion protein to assess parkin ubiquitination activity [39]. Both parkin and pParkin can efficiently poly-ubiquitinate Ypet_TEV_-pUb (Fig. 10, lanes 1-3). Following TEV cleavage, only a single pUb band was visible, indicating parkin did not create chains on the pUb anchor (Fig 10, lane 5). Similar to the shift in mG2~Ub_n_, a downward shift in the YPet~Ub_n_ laddering was also observed (Fig. 10, lane 6) indicating that parkin ubiquitinates lysine residues on the YPet protein. These results confirm a function of pUb is to act as the substrate for parkin and anchor an OMM protein for ubiquitination.

**Figure 10:**
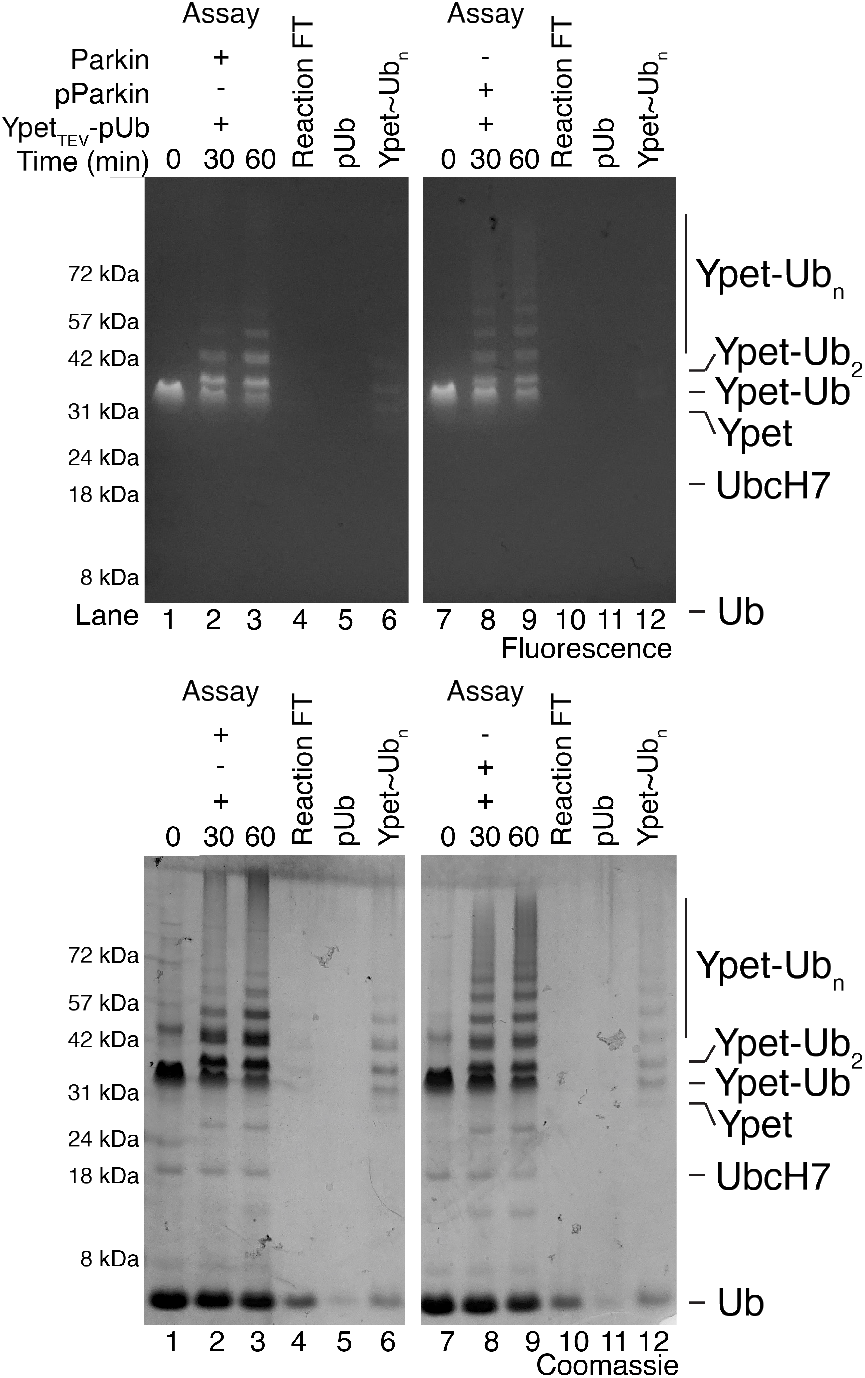
Parkin ubiquitinates an unnatural acceptor that is linked to pUb. Ubiquitination assay using Ypet_TEV_--pUb. The reactions and purifications were resolved using 12% Bis-Tris SDS-PAGE and visualized using YPet fluorescence on a BluPAD LED Transilluminator (FroggaBio) (top) or by Coomasssie staining (bottom).

## Discussion

Understanding the mechanisms of activation and substrate ubiquitination by the E3 ligase parkin are key to identifying its role in early-onset PD and the development of potential therapeutics. Current models show parkin recruitment to damaged mitochondria by PINK1-phosphorylated pUb [31, 39]. These pre-existing ~pUb molecules likely arise from constitutive ubiquitin signaling mediated by a number of constitutive mitochondrial E3 ligases [39, 40, 54], followed by PINK1-mediated phosphorylation of surface-exposed ubiquitin, triggered by oxidative stress. Parkin can then be phosphorylated itself by PINK1, creating a pParkin/pUb complex that in autoubiquitination assays demonstrates maximal ubiquitination activity [8, 17, 30]. Indeed, this step wise activation of parkin activity has been implicated numerous times with both *in vivo* and *in vitro* systems. Yet this model reveals several unresolved gaps in the mechanism. First, while numerous OMM proteins are shown to be ubiquitinated by parkin under oxidative stress conditions [22, 26], the only common feature across these targets is their location on the OMM. OMM target proteins such as Miro1, CISD1, mitofusins 1/2 and proteins in the TOM complex have no identified sequence similarity to suggest a common parkin recognition motif for a targeted lysine residue. Further, there is little information to suggest direct interaction of these proteins with parkin, a requirement for ubiquitination. Second, in most ubiquitination assays parkin shows a preference to ubiquitinate itself, regardless of the target protein present in an assay, especially with fully activated pParkin/pUb. This is an odd phenomenon for an enzyme, especially for an E3 ligase that is targeting proteins for signaling or degradation.

Our work shows that the interaction of Miro1 or CISD1 with parkin is only observed when a pUb molecule is tethered to either acceptor protein. We could not detect a direct interaction between parkin and these two putative substrates, even when these proteins are ubiquitinated. Further, our data shows that ubiquitination of Miro1 or CISD1 is very poor compared to autoubiquiti nation of parkin even in the pParkin/pUb state. The interaction of an acceptor-pUb is moderately weaker than that of parkin with pUb alone [8, 17] but establishes that pUb is the main determinant for parkin recognition that brings an acceptor protein nearby to facilitate ubiquitination. This observation that pUb is the “substrate” for parkin reconciles the broad spectrum of proteins targeted for ubiquitination by the E3 ligase. Multiple studies have identified more than 30 OMM proteins as targets for parkin ubiquitination. In addition, comparable numbers of non-mitochondrial proteins in the cytoplasm, endoplasmic reticulum and nucleus are also ubiquitinated by parkin [22, 26] where it is unclear how phosphorylation would regulate the process. This could suggest that pUb or a similar molecule exists outside the OMM that is available for parkin binding and activation. One possibility is the ubiquitin-like protein SUMO-1, demonstrated to bind and localize parkin to the nucleus [57]. Further, parkin with an N-terminal yeast SUMO-tag shows higher ubiquitination activity than parkin alone [6, 42, 43] perhaps indicating that the SUMO family of modifiers are all able to replace pUb as an activator. Consistent with this, SUMOs have 3-4 Asp/Glu residues that structurally align with the S65 loop in ubiquitin and could mimic phosphate binding as observed with pSer65 [12]. A similar Ub-like molecule, ISG15, was also observed to aid in parkin translocation [58]. If other ubiquitin-like molecules can bind and activate this intermediate level of parkin function, then it explains parkin substrates beyond the mitochondrion and expands parkin’s role in cellular function.

We observed robust ubiquitination of mG2-pUb or CISD1-pUb proteins by parkin that occurred on the acceptor moiety rather than pUb. This indicates that pUb only acts to guide and orient the acceptor protein to parkin. Previous work by Okatsu et al. (2015) had suggested the pUb is a genuine substrate for parkin and our work supports this idea. Moreover, our cleavage assays demonstrated that parkin ubiquitinates the tagged protein and not pUb, similar to that observed in a previous study [39], but also that parkin only requires surface-exposed lysines and can ubiquitinate even unnatural substrates as we demonstrated with Ypet. Intriguingly, other studies have noted that a large N-terminal-tag on parkin (MBP) was also exclusively ubiquitinated by parkin further supporting the requirement of parkin-substrate tethering [6, 27]. We also showed that an acceptor-pUb retains binding to the basic patch containing H302 [32, 39, 59]. These findings rationalize immunoprecipitation assays from cell lysates that have hinted at interactions between parkin and some substrates including Miro1 [35, 36] and mitofusin 2 [37] that are enhanced after oxidative stress, likely due to pUb tagging of each of these proteins. The commonality of pUb on many different OMM proteins provides a biological explanation for the breadth of parkin substrates.

The stimulation of acceptor-pUb ubiquitination requires that an E2~Ub conjugate can interact with parkin and that an increase in access to the catalytic cysteine (C431) occurs. In the auto-inhibited state, the Ubl domain sits between the RING1 central helix and the IBR domain [8, 17]. Multiple three-dimensional structures show that upon pUb binding to the RING0-RING1 cleft, straightening of helix H3 moves the IBR domain away from the Ubl domain [33, 45, 46]. Several other movements have been noted that make subtle alterations to other intradomain interactions. This results in weaker Ubl association in parkin/pUb compared to the auto-inhibited state [8, 17]. This step also improves the efficiency of Ubl domain phosphorylation [31]. Notably a significant pocket between the Ubl-RING1-IBR domains is liberated that easily fits a ubiquitin molecule based on structural and biochemical experiments [33]. Recent structures show that UbcH7 and UbcH7-Ub interact with regions of the parkin RING1 domain (L1, L2 loops) [44, 46] that are minimally involved in intermolecular contacts with the Ubl domain (Fig. 11). This structure also shows the conjugated ubiquitin is mobile but can occupy the Ubl-RING1 pocket with minimal interference of the Ubl domain. Therefore, it is conceivable that the widened RING1/IBR binding pocket could accommodate both Ubl and UbcH7-Ub. Consistent with these observations, isothermal titration experiments show dissociation constants show moderate binding for parkin/pUb with UbcH7~Ub (K_d_ ~40 *μ*M, unpublished) and affinity experiments demonstrate that UbcH7 can pull-down full-length parkin [6]. Further, NMR based titration studies demonstrated an interaction between parkin:pUb and UbcH7 that was enhanced by introduction of the parkin activating mutation W403A within the REP element of the tether [17, 44], a finding that is functionally supported by *in organello* ubiquitination assays which observed increased mitofusin 2 ubiquitination with parkin^W403A^ compared to WT [59]. This suggests that UbcH7 binding relies more on displacement of the tether on RING1 and not on full release of the Ubl domain by phosphorylation.

**Figure 11:**
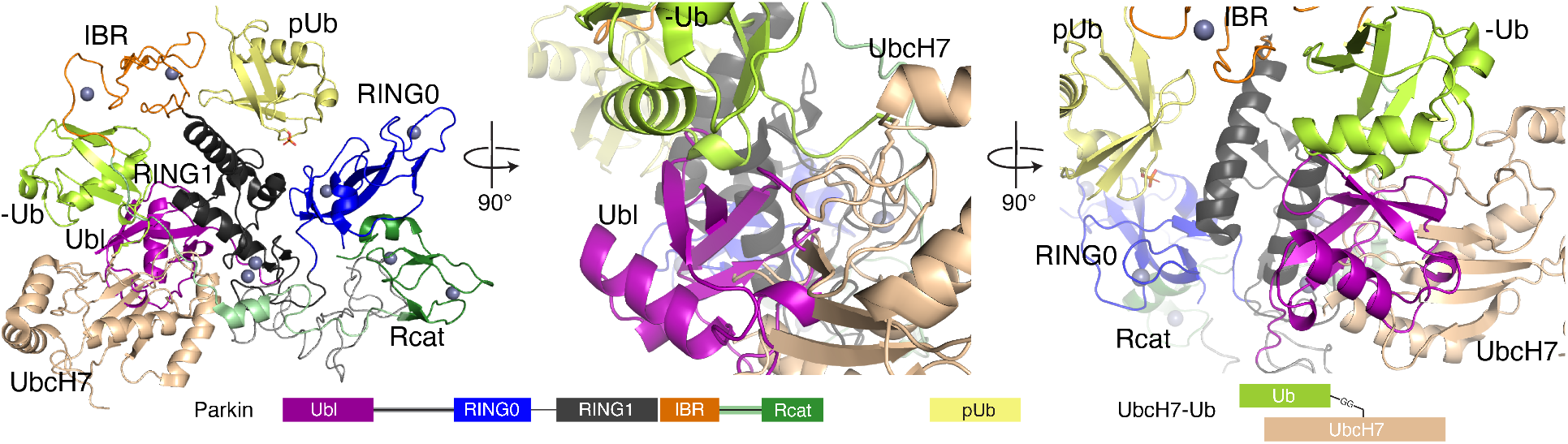
UbcH7-Ub and Ubl could simultaneously bind parkin:pUb. Three views of parkin:pUb bound to UbcH7-Ub. A model of R0RBR:pUb:UbcH7-Ub (PDB 6N13) was modified by adding a modelled Ubl-linker (aligned from PDB: 5N2W). The linker was modeled by Modeller [55] in Chimera [56] and the structures were aligned and imaged using PyMOL.

Our novel observations indicate that acceptor-pUb species are multi-ubiquitinated by parkin, in the absence of phosphorylation of its Ubl domain and that subsequent phosphorylation results in enhanced parkin autoubiquitination. This divergent mechanism accelerates ubiquitination of mitochondrial proteins upon initial recruitment of parkin to the OMM, without relying on phosphorylation of the Ubl domain (Fig. 12A). Instead, this mechanistic step ensures that additional Ub molecules are placed on target proteins and are available for PINK1 phosphorylation and subsequent parkin recruitment, contributing to a feed-forward mechanism of parkin mediated ubiquitination [31, 39]. After initial recruitment to the OMM, gradually parkin will also become phosphorylated by PINK1 which in turn, increases parkin activity towards autoubiquitination (Fig. 12B). This mechanism agrees with experiments in *Drosophila melanogastor* expressing unphosphorylatable parkin (S65A) that have a partially rescued phenotype compared to parkin-null flies [10] and parkin^S65A^ co-localizes to TOM20 on the mitochondrial membrane as efficiently as wildtype parkin [60]. This would create a level of mitochondrial ubiquitination sufficient for mitophagy signaling before parkin itself becomes phosphorylated and triggering parkin autoubiquitination. However, the direct consequence of parkin autoubiquitination is not clear and how ubiquitination of the E3 ligase alters its recruitment of the E2~Ub conjugate or catalytic activity.

**Figure 12:**
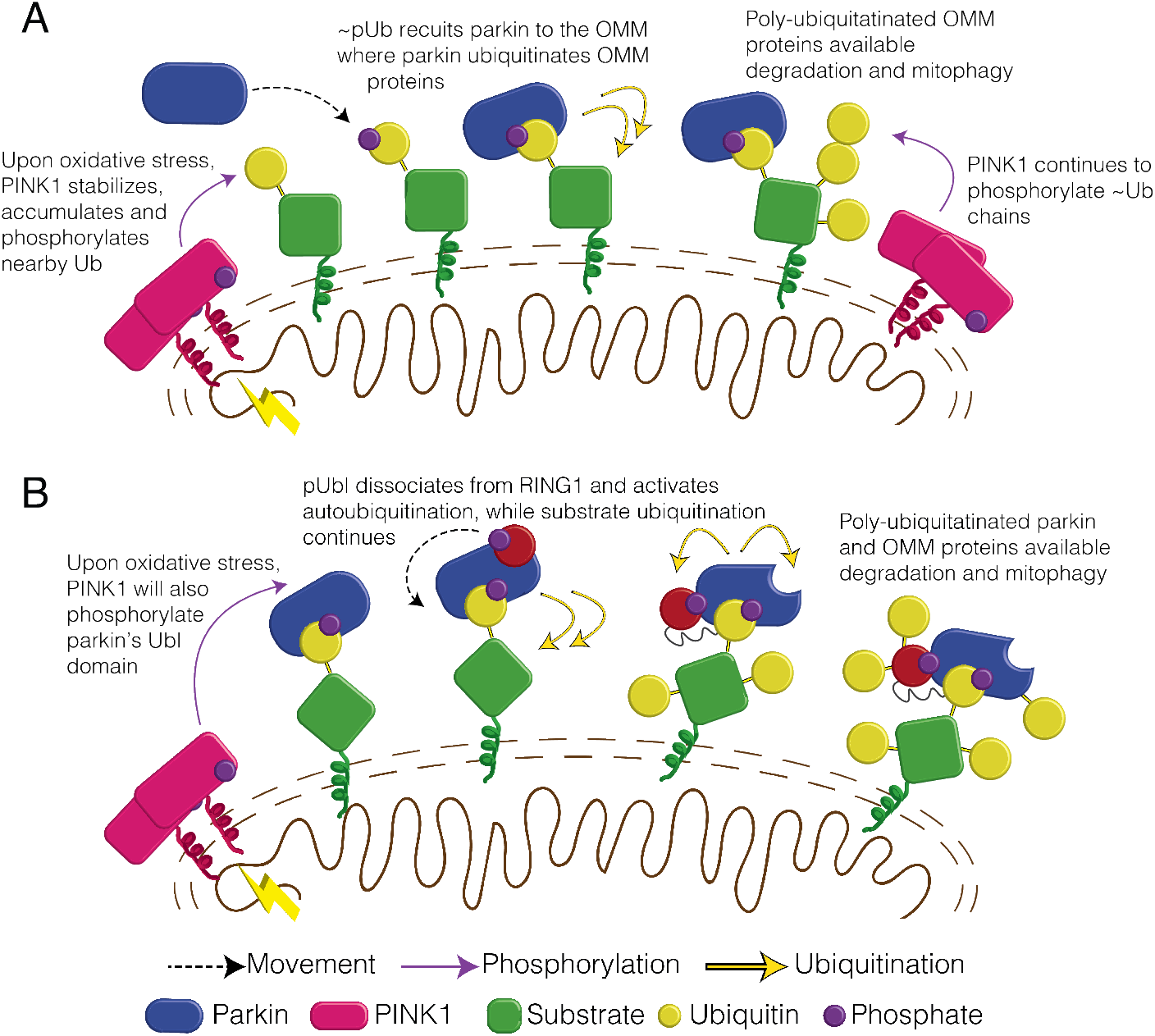
Schematic of divergent parkin ubiquitination activity. (*A*) Parkin is recruited to mitochondria by surface pUb molecules and ubiquitinates OMM proteins. These new -Ub molecules are phosphorylated, thus amplifying the signal. (*B*) Parkin’s presence at the OMM will also lead to Ubl phosphorylation. This also increases parkin’s ubiquitination activity and leads to autoubiquitination.

Over 70 substitutions in parkin have been associated with early-onset PD. Some of those mutations are in regions of parkin that existing mechanistic knowledge has made clear how parkin function is affected, for example E2~Ub binding at RING1 (T240R) [44], loss of the catalytic cysteine (C431F) [16], loss of the proposed pUbl rebinding site (K161N, K211N) [16, 43, 46] or even substitutions throughout the Ubl domain differentially affecting the rate of parkin phosphorylation [8, 16, 61, 62]. However, the impact on parkin function for many of the PD-related substitutions is not as predictable and the available parkin structures are insufficient to identify which part of the mechanism is affected, Strikingly, none of the known substitutions appear to have a detrimental effect directly on pUb binding at the phosphate binding basic patch. The novel observations in this work will be important for influencing further structural and functional studies for elucidating parkin’s mechanism of substrate ubiquitination prior to phosphorylation. Any useful therapeutic intervention targeting parkin will be dependent on an understanding of the consequences of altered Parkin activity. For example, if parkin activators are developed with therapeutic intentions, knowledge of the impact of such activators on parkin levels, parkin activity, and substrate targeting will be key to establishing appropriate paradigms. Our study offers a clear insight into the effects of parkin modulation, by activators and modifiers, on substrate choice, and thus has implications for the rational design of therapeutics for mitochondrial targeting.

## Acknowledgements

This research was supported by a grant (PJT 166019) from the Canadian Institutes of Health Research (GSS) and a grant (209347/Z/17/Z) from Wellcome Trust UK (HW). KMD was the recipient of a Natural Sciences and Engineering Research Council postgraduate scholarship.

## Author Contributions

KMD conceived the study, designed experiments, completed protein interaction and ubiquitination experiments, analyzed data and wrote the manuscript. ACR-D conducted protein interaction and ubiquitination assays for CISD1, analyzed data and wrote the manuscript. GS and RT conducted thermophoresis experiments and analyzed data. DH synthesized and purified proteins and conducted ubiquitination assays. KRB synthesized and purified proteins. HW designed experiments, analyzed data and wrote the manuscript. GSS conceived the study, designed experiments, analyzed data and wrote the manuscript.

## Conflict of Interest

The authors declare that they have no conflicts of interest.

## Notes

### Competing Interest Statement

The authors have declared no competing interest.

