## Supplementary material for "Distinct phosphorylation signals drive acceptor versus self-ubiquitination selection by Parkin": Fig. S1

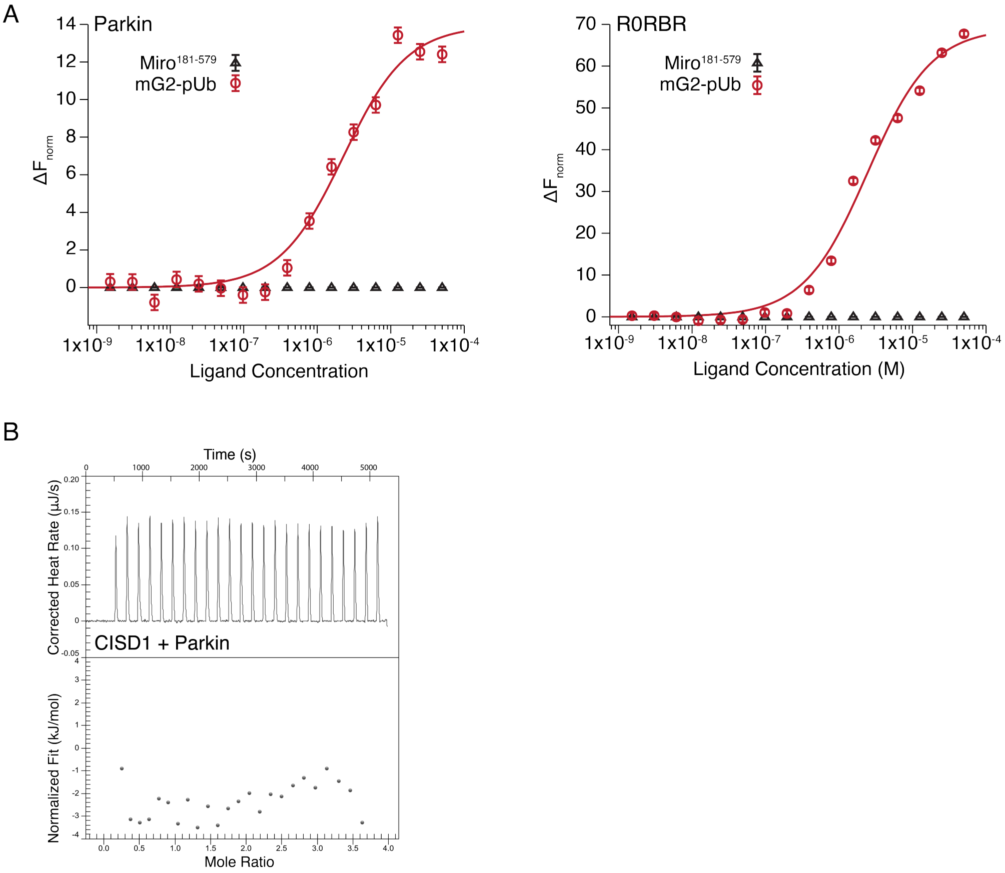


Figure S1: Parkin does not interact with Miro1$^{181\text{-}579}$ or CISD1. (*A*) MST experiment titrating 16 dilutions of Miro1$^{181\text{-}579}$ (black triangle) or mG2-pUb (red circles) into $His_6$-Smt3-tagged parkin (left) and R0RBR (right). (*B*) Isothermal titration calorimetry of parkin (20 $\mu$M) titrated with CISD1 (200 $\mu$M).
