## Supplementary material for "Distinct phosphorylation signals drive acceptor versus self-ubiquitination selection by Parkin": Fig. S2

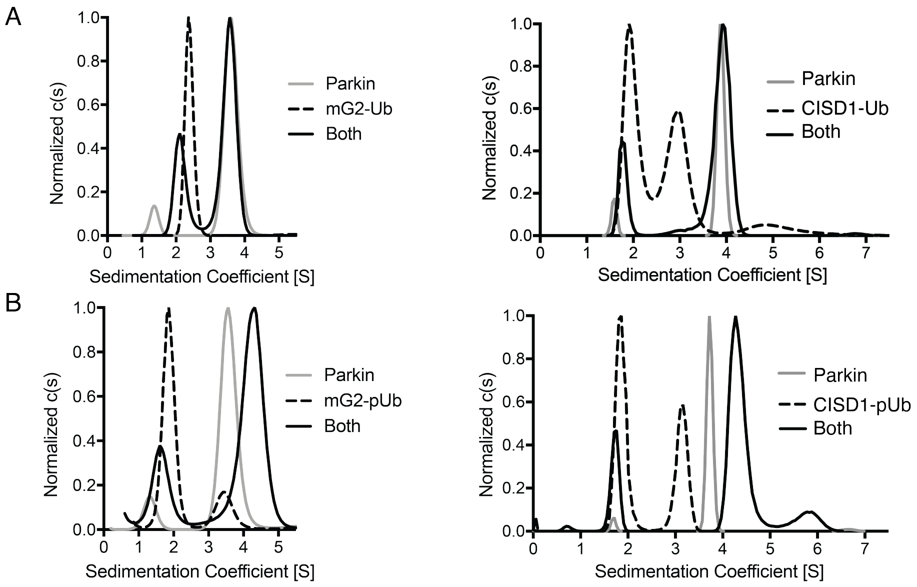


Figure S2: Parkin interacts with mG2-pUb and CISD1-pUb. For mitochondrial acceptors (*A*) mG2 and (ltextitB) CISD1, sedimentation velocity experiments were conducted to determine the sedimentation of parkin alone (grey), substrate-(p)Ub alone (dashed) and parkin/substrate-(p)Ub (black).
