## Supplementary material for "Distinct phosphorylation signals drive acceptor versus self-ubiquitination selection by Parkin": Fig. S3

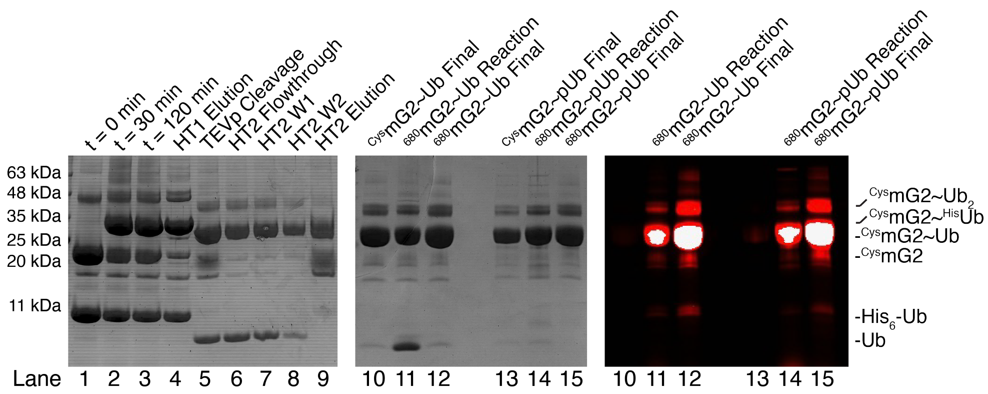


Figure S3: Preparation of monoubiquitinated mG2, mG2$^{K572}$~Ub. Preparative ubiquitination reaction (lanes 1-3), purification (lanes 4-9) and fluorescent labeling (lanes 10-21) of mono-ubiquitinated mG2$^{K572}$~Ub and mG2$^{K572}$~pUb.
